# Salt stress reverses root circumnutation, -skewing and -growth direction in Arabidopsis

**DOI:** 10.64898/2026.08.27.747533

**Authors:** Hui Sheng, Ringo van Wijk, Harro J. Bouwmeester, Teun Munnik

## Abstract

Plant roots exhibit remarkable developmental plasticity, resulting in the adaptation of growth direction and architecture upon environmental changes. Previously, we demonstrated that inorganic phosphate (P_i_) triggers Arabidopsis roots to skew to the left when grown on tilted agar plates. This so-called ’phosphate-dependent skewing’ (PDS) is caused by a right-handed (clockwise, CW) circumnutation of the root tip, which is driven by a left-handed (counterclockwise, CCW) cell file rotation (CFR) of epidermal cells in the root elongation zone, and involves the cortical microtubule cytoskeleton (Sheng et al., 2024). In the present study, we demonstrate that NaCl triggers a skewing response in the opposite direction and that all other helical movements are also reversed. Thus, ’Salt-Induced Rightward Skewing’ (SIRS) is accompanied by a right-handed (CW) epidermal CFR, a left-handed (CCW) circumnutation of the root tip, and hence, a left-handed (CCW) helical root growth. Comparing different Na^+^- and Cl^-^ salts revealed that SIRS is predominantly caused by cations, and can be induced by K^+^ and osmotic stress as well, although Na^+^ is most efficient. To get further insight into the mechanism underlying this response, we tested candidate genes from an earlier GWAS on root responses to salt stress (Deolu-Ajayi *et al*., 2019) for their potential involvement. This identified *GLT1* and *DOB1* as being involved in the root skewing response to P_i_ and NaCl, respectively. Our findings reveal that P_i_ and salinity elicit opposing effects on root circumnutation, and hence root skewing and growth direction. Understanding the molecular machinery driving this helical behaviour may help explain adaptive mechanisms, including changes in the spatial architecture of roots, and may facilitate the optimization of crop yield under abiotic stress conditions through breeding or crop management strategies. Our results also shed new light on halotropism, which is typically measured as a change in root growth direction to the right, which in the present study has been identified to represent SIRS.

## Introduction

Plant roots facilitate uptake, storage, and translocation of nutrients and water, are engaged in communication and interaction with the microbiome and neighbouring plants, and are responsible for anchoring plants into the soil (Maurel & Nacry, 2020; Karlova *et al*., 2021). The developmental plasticity of roots is very high, allowing them to detect environmental changes and modify their architecture and growth direction to optimize water- and nutrient acquisition while avoiding obstacles and toxic compounds (Shahzad & Amtmann, 2017; Gupta *et al*., 2020; Zou *et al*., 2022; González-García *et al*., 2023; Voothuluru *et al*., 2024; Zhang *et al*., 2024). Understanding such adaptive mechanisms, and unraveling the molecular basis of such response programs, are of vital importance to secure crop yield in the face of increasing environmental stress challenges as a result of climate change (Lynch, 1995; Munns & Tester, 2008; Munns *et al*., 2020; Shelden & Munns, 2023).

In his famous book, *"The Power of Movements in Plants"*, Charles Darwin already categorized plant movements into tropisms, nastic movements, and nutations (Darwin & Darwin, 1880), which often occur simultaneously. For example, the distinctive movements of Arabidopsis roots on tilted agar plates, like skewing, coiling, and waving, are believed to arise from an interplay between circumnutation, gravitropism, and thigmotropism (Migliaccio & Piconese, 2001; Oliva & Dunand, 2007; Roy & Bassham, 2014; Shih *et al*., 2014; Swarbreck *et al*., 2019; Sipos & Várkonyi, 2022; Sheng *et al*., 2024; Porat *et al*., 2024). Root skewing is marked by the characteristic twisting of epidermal cell files, known as cell file rotation (CFR), where left-handed (counterclockwise; CCW) epidermal CFR corresponds to leftward skewing when seen from the front of the plate, and *vice versa* (Rutherford & Masson, 1996; Furutani *et al*., 2000; Yuen *et al*., 2005). Both skewing and CFR were demonstrated to be associated with the microtubule cytoskeleton through the use of mutants (Thitamadee *et al*., 2002; Nakajima *et al*., 2004; Shoji *et al*., 2004; Molines *et al*., 2018; Buschmann & Borchers, 2020), as well as pharmacological drugs affecting microtubule polymerization (Oliva & Dunand, 2007; Hodge *et al*., 2009). Recently, we demonstrated that the root skewing of Arabidopsis seedlings on vertical ½MS-agar plates is caused by inorganic phosphate (P_i_) in the medium, hence coined ’phosphate-dependent skewing’ (PDS). P_i_ promotes leftward root skewing and right-handed (clockwise, CW), circumnutation, which is accompanied by a left-handed (CCW) epidermal CFR, and a reorganisation of the cortical microtubule cytoskeleton in the root elongation zone (Sheng *et al*., 2024), providing unprecedented insight into the role of P_i_ in directional root growth, microtubule responses and circumnutation.

Salinity is an increasing threat to modern agriculture, affecting over 800 million hectares of land worldwide, including more than 20% of irrigated areas (Ghassemi *et al*., 1995; Munns & Tester, 2008; Acosta-Motos *et al*., 2017; Hasanuzzaman & Fujita, 2022; Negacz *et al*., 2022). Various physiological processes are negatively affected by salinity, including photosynthesis, cell growth and division, and stomatal conductance (Dzinyela *et al*., 2023). Salinity comprises both osmotic (water deficit) and ionic (accumulation of Na^+^ and Cl^-^) stress components (Munns & Tester, 2008; Wu, 2018; van Zelm *et al*., 2020; Su *et al*., 2021). Osmotic stress occurs first, through the reduction of the external water potential, impairing the plant’s ability to take-up water (Farooq *et al*., 2015). Simultaneously, Na^+^ ions enter cells through non-specific cation channels, which lowers the cytoplasmic water potential resulting in excessive water-influx and subsequent swelling of the cells (Colin *et al*., 2022; Dzinyela *et al*., 2023). The increase of Na^+^ in cells and tissues also causes ionic stress, especially in green tissues where it interferes with photosynthesis, which has more serious consequences than the osmotic stress (Chowaniec & Rola, 2022). Influx of Na^+^ also causes depolarisation of the membrane potential, which results in extrusion of K^+^ (Assaha *et al*., 2017). As essential metal co-factors of various metabolic enzymes, the increase of Na^+^ and decrease in K^+^ can have dramatic effects on both primary- and secondary metabolism (Dzinyela *et al*., 2023). Excess of Na^+^ may also displace Ca^2+^ from particular binding sites, which in cell walls reduces the crosslinking of pectin, weakening wall structure and affecting cell elongation (Proseus & Boyer, 2012; Feng *et al*., 2018; Byrt *et al*., 2018).

In the present study, the effect of salt stress on PDS was analysed. Surprisingly, we found that NaCl reverses the skewing direction of PDS and that all helical movements involved, were also reversed. Testing candidate genes from a recent GWAS analysis to early salt stress, using T-DNA insertion mutants, the potential involvement of *GLT1* and *DOB1* under P_i_ and NaCl conditions, respectively, were identified. Our results shed new light on the helical growth of roots and the effects of NaCl on this process.

## Materials and Methods

### Plant material & Growth

*Arabidopsis thaliana* accessions Columbia (Col-0) and Wassilewskija (Ws-4) were used. T-DNA insertion lines (all Col-0) include (AT5G53460) *glt1-1* (SALK_115735C), *glt1-2* (SALK_092158C), *glt1-4* (SALK_033098) and *glt1-5* (SALK_072454); (AT3G60140) *din2-1* (SALK_029737C) and *din2-2* (SALK_069359); and (AT4G25670) *dob1-1* (SALK_056459C) and *dob1-2* (SALK_203487C), which were obtained from NASC (Nottingham Arabidopsis Stock Centre). Additional T-DNA lines tested are listed in Supplementary Table S1. Seeds were sterilized with 1 mL 50% bleach and 60 μL 70% ethanol for 10 min and then washed 5 times with sterile water. Sterilized seeds were sown on square Petri dishes (12 ×12 cm) containing 40 ml of ½ strength Murashige-Skoog (½MS) medium (Murashige & Skoog, 1962), supplemented with 1% (w/v) sucrose and 0.8% (w/v) Micro agar (Duchefa), and with pH adjusted to 5.8 with KOH. After stratification at 4 °C for 2 days in the dark, agar plates were placed vertically (70° angle) in a growth chamber at 22 °C (16-hour light / 8-hour dark cycle).

To investigate root growth behaviour at different NaCl- and P_i_ concentrations, ½MS medium *without* phosphate (Duchefa) was used, which was supplemented with P_i_ from a 100 mM KH_2_PO_4_ stock and NaCl from 1 M stock. Differences in K^+^ were compensated with KCl (Sheng *et al*., 2024). Normally, ½MS contains 625 μM P_i_ but to have a slightly stronger PDS response in Col-0, 800 μM P_i_ was often used. To study the effect of NaCl and validate the effect of P_i_, low (300 μM) and high (2500 μM) P_i_ concentrations were used.

### Quantification of root skewing

Agar plates were scanned from the back using an Epson Perfection V800 Photo Scanner (J221B). Images were then flipped horizontally to simulate a front-view orientation and to see the real skewing direction. Root length (L), angle of deviation from the gravity of the root tip (θ), and length of the idealized root response (L_c_) were measured using ImageJ to calculate the horizontal growth index (HGI), as described earlier (Sheng *et al*., 2024).

### Root growth direction on horizontal agar plates

Sterilized seeds were sown on ½MS medium containing low or high P_i_ (300 and 2500 μM) ± 50 mM NaCl, and grown vertically (70° angle) for 4 days. Root tips were then marked on the back of the plate by pen, and the plates placed horizontally. After two days, plates were scanned to visualize the root growth direction. Plates were scanned again after 6 or 10 days to visualize lateral root growth direction.

### Epidermal cell file rotation analysis

A Leica stereomicroscope (Leica MZFLIII, Leica Microsystems GmbH, Wetzlar, Germany) was used to visualize the root surface of Arabidopsis seedlings. Angles of the epidermal CFR were measured as described previously (Sheng *et al*., 2024). To indicate the two different rotational directions, left-handed, counter-clockwise CFR was designated as positive, while right-handed, clockwise CFR as negative (Suppl. Fig. S1a).

### Genotyping T-DNA insertion lines

Seeds of T-DNA insertion lines were propagated in soil and grown for 3-4 weeks in a growth chamber at 22 °C (12-hour light / 12-hour dark cycle). Leaf material was pressed on a Whatman^®^ FTA card (Sigma-Aldrich). Discs (1.2 mm), containing DNA from the samples, were punched out and deposited into PCR tubes. Discs were washed with 50 μL FTA buffer (10 mM Tris pH 7.5, 2 mM EDTA, and 0.1% (v/v) Tween-20), and 150 μL with TE^-1^ buffer (10 mM Tris pH 8.0, 0.1 mM EDTA) twice, 5 min each. PCR reaction buffers and genotyping primers (Suppl. Table S2) were added and PCR started, with a program of 20 sec denaturation at 95°C, 30 sec annealing at 54°C, and 90 sec elongation at 72°C, for 35 cycles. Resulting DNA samples were analysed by agarose gel electrophoresis to confirm T-DNA insertions and to select homozygous plants.

### qRT-PCR analysis

RNA from roots of 9-day-old seedlings was isolated using the CTAB method. Samples were treated with TURBO^TM^ DNase (Invitrogen), and cDNA was synthesized from 1 μg of total RNA using reverse transcriptase (Fermentas). Expression levels of *GLT1* (At5g53460), *ACBP1* (At5g53470), *DIN2* (AT3G60140) and *DOB1* (AT4G25670) were measured using qRT-PCR with Eva-Green kit (Bio-connect) using *SAND* (At2g28390) as reference gene. RT-PCR primers are listed in Table S2.

### Statistical analysis

Statistical analyses were performed using GraphPad Prism 10. The Shapiro-Wilk test was applied to analyze the normal distribution of values. One-way or two-way analysis of variance (ANOVA) followed by Tukey’s or Šídák’s multiple comparison tests was used. If values were not normally distributed, the Kruskal-Wallis test followed by Dunn’s multiple comparison tests were used. Different letters indicate significant differences between different treatments (p<0.05). Asterisks indicate a statistically significant difference among different treatments. Significantly different means (* p<0.05, ** p< 0.01, *** p< 0.001; ns, not significant) were separated by Tukey’s NEJM.

## Results

### Salt stress reverses the skewing direction of roots

Previously, we demonstrated that the typical skewing response of Arabidopsis roots on tilted ½MS-agar plates is caused by the P_i_ present in the medium. P_i_ dose-dependently promotes a rightward, clockwise (CW) rotational movement (circumnuation) of the root tip, which during growth on plates manifests itself as a leftward skewing response, hence coined ’Phosphate-Dependent Skewing’ (PDS) (Sheng *et al*., 2024). Since salt stress has strong effects on root growth and architecture (Wang *et al*., 2009; Galvan-Ampudia *et al*., 2013; Julkowska *et al*., 2014), we decided to investigate the effect of NaCl on PDS. Accordingly, Arabidopsis seedlings (Col-0 and Ws-4) were grown on vertical (70°) ½MS agar plates containing 800 μM P_i_ KH_2_PO_4_, slightly more than the 625 μM normally present in ½MS medium, to trigger a stronger PDS response. The medium was supplemented with various concentrations of NaCl, ranging from 0 to 125 mM. In the absence of NaCl, seedlings skewed to the left, with Ws-4 typically showing a much stronger response than the Col-0 ecotype, as demonstrated previously (Sheng *et al*., 2024). With NaCl, however, a dose-dependent reversion of the skewing direction from leftward to rightward was observed. In addition, salt stress also clearly inhibited root growth, with Col-0 being slightly more sensitive (see 75 mM; Fig. 1; Suppl. Fig. S1b).

**Fig. 1.**
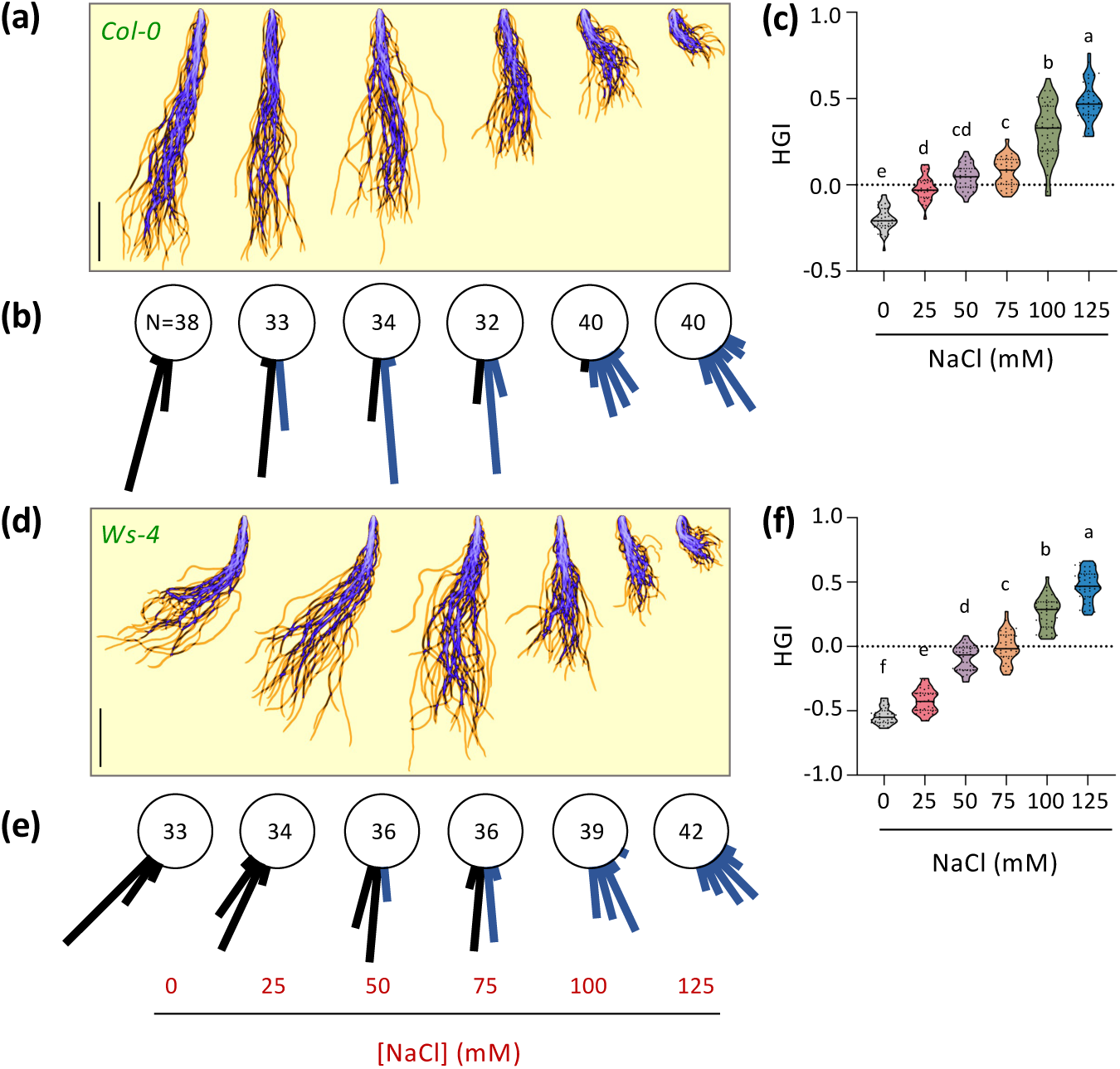
NaCl promotes rightward root skewing in Arabidopsis seedlings. Arabidopsis ecotype Col-0 (a-c) and Ws-4 (d-f) were grown on ½MS-agar medium containing 800 µM KH_2_PO_4_ and different concentrations of NaCl (0-125 mM). After 9 days, roots were scanned and their Horizontal Growth Index (HGI) determined. Without NaCl, roots skew to the left due to P_i_, a process named Phosphate Dependent Skewing (PDS). NaCl reverses this and makes roots skew the opposite direction, to the right. (a, d) Primary root projections with the overlap indicated in purple. (b, e) Root angles (in 10° categories) with the number of roots measured indicated. 0° equals vertical root growth. Black and blue lines represent negative (-) and positive (+) root angles, and left- and rightward skewing directions, respectively. (c, f) HGI values. The middle line represents the median, the dotted line represents quartiles and individual data points are represented by dots. Different letters indicate significant differences (*p*<0.05) among treatments using One-way ANOVA followed by Tukey HSD post-hoc analysis: *F*_5,211_ = 210.3, *p* < 0.001 (B) or *F*_5,214_ = 511.9, *p* < 0.001 (D). Scale bar = 1 cm.

### Effect of phosphate and NaCl on root skewing

The rightward skewing response in the presence of NaCl is the opposite of the skewing direction induced by P_i_, PDS. To further investigate the individual contributions of NaCl and P_i_, the effect of three NaCl concentrations (0, 50, and 100 mM) in combination with four different P_i_ concentrations (300, 625, 1250, and 2500 μM KH_2_PO_4_) was investigated (Fig. 2). Without NaCl, the leftward PDS response is obvious in both Col-0 (Fig. 2a) and Ws-4 (Fig. 2b). However, with 50 mM NaCl, root skewing was completely reversed to the right for Col-0 (Fig. 2c) while causing a straightening of the roots in Ws-4 (Fig. 2d). At 100 mM NaCl, both Col-0 and Ws-4 (Fig. 2e, 2f) skewed to the right, even though growth was significantly inhibited.

**Fig. 2.**
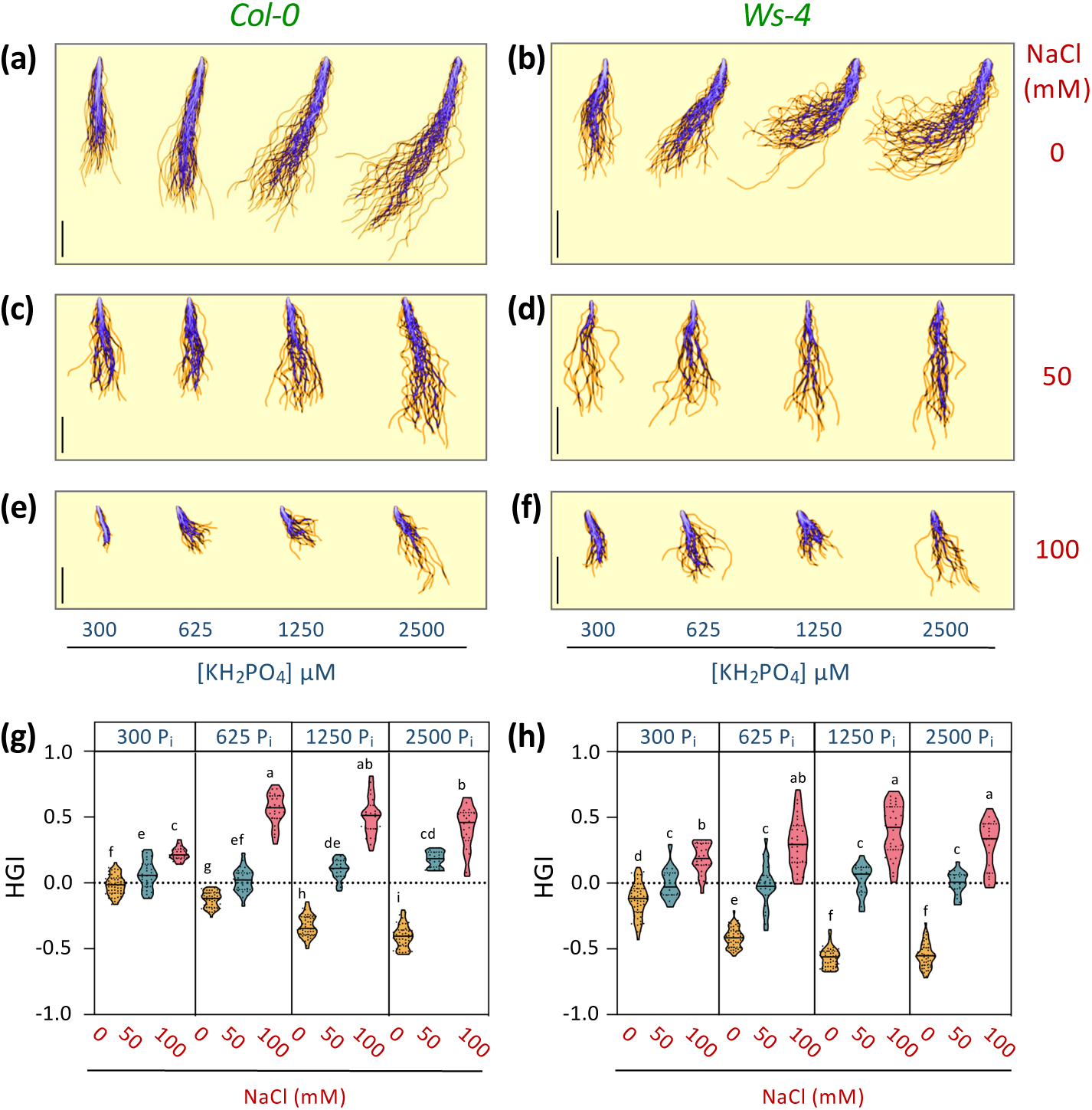
Effect of NaCl and P_i_ on PDS in Arabidopsis Col-0 and Ws-4 seedlings. (a, b) Projections of 9-day-old seedlings of Col-0 (A) and Ws-4 (b) grown on ½MS medium with different KH_2_PO_4_ concentrations. (c-f) Projection of primary roots of Col-0 (c, e) and Ws-4 (d, f) seedlings grown at the indicated P_i_ concentrations on a medium with 50 mM (c, d) or 100 mM (e, f) NaCl. (g, h) HGI for Col-0 (g) and Ws-4 (h). The middle line represents the median, the dotted line represents quartiles and individual data points are represented by dots. Two-way ANOVA was performed to identify significant differences between treatments. Different letters indicate significant differences (*p*<0.05) among treatments by Tukey HSD post-hoc analysis between different P_i_ concentrations of (g) (*F*_2,379_ = 1508, *p* < 0.001) and (h) (*F*_2,363_ = 1046, *p* < 0.001); the significance between NaCl treatments of (g) (*F*_2,379_ = 13,18, *p* < 0.001) and (h) (*F*_3,363_ = 8,356, *p* < 0.001). N = 15-52. Scale bar = 1 cm.

To assess the impact of P_i_ on the salt-induced root skewing, HGI values at different NaCl concentrations were plotted per P_i_ concentration (Fig. 2g-h). In general, each P_i_ concentration showed the same trend, with most significant differences noted between the lowest- and highest P_i_ concentration tested (i.e. 300 and 2500 μM).

### NaCl-induced skewing is predominantly ionic

To investigate whether the NaCl induced-skewing response is ionic- or osmotic in nature, ½MS-agar plates with 800 μM KH_2_PO_4_ were supplemented with 50 mM NaCl or 100 mM mannitol, which has the same osmotic strength (osmolality). NaCl displayed a much stronger effect on root skewing than mannitol in both Col-0 and Ws-4 seedlings (Fig. 3a, b). Mannitol did have an effect on PDS in Col-0, reducing the leftward skewing induced by P_i_ and causing the roots to straighten. This did not occur in Ws-4, probably because the PDS is much stronger in this ecotype (Sheng et al., 2024). In both ecotypes, mannitol had a stronger negative impact on root growth than NaCl.

**Fig. 3.**
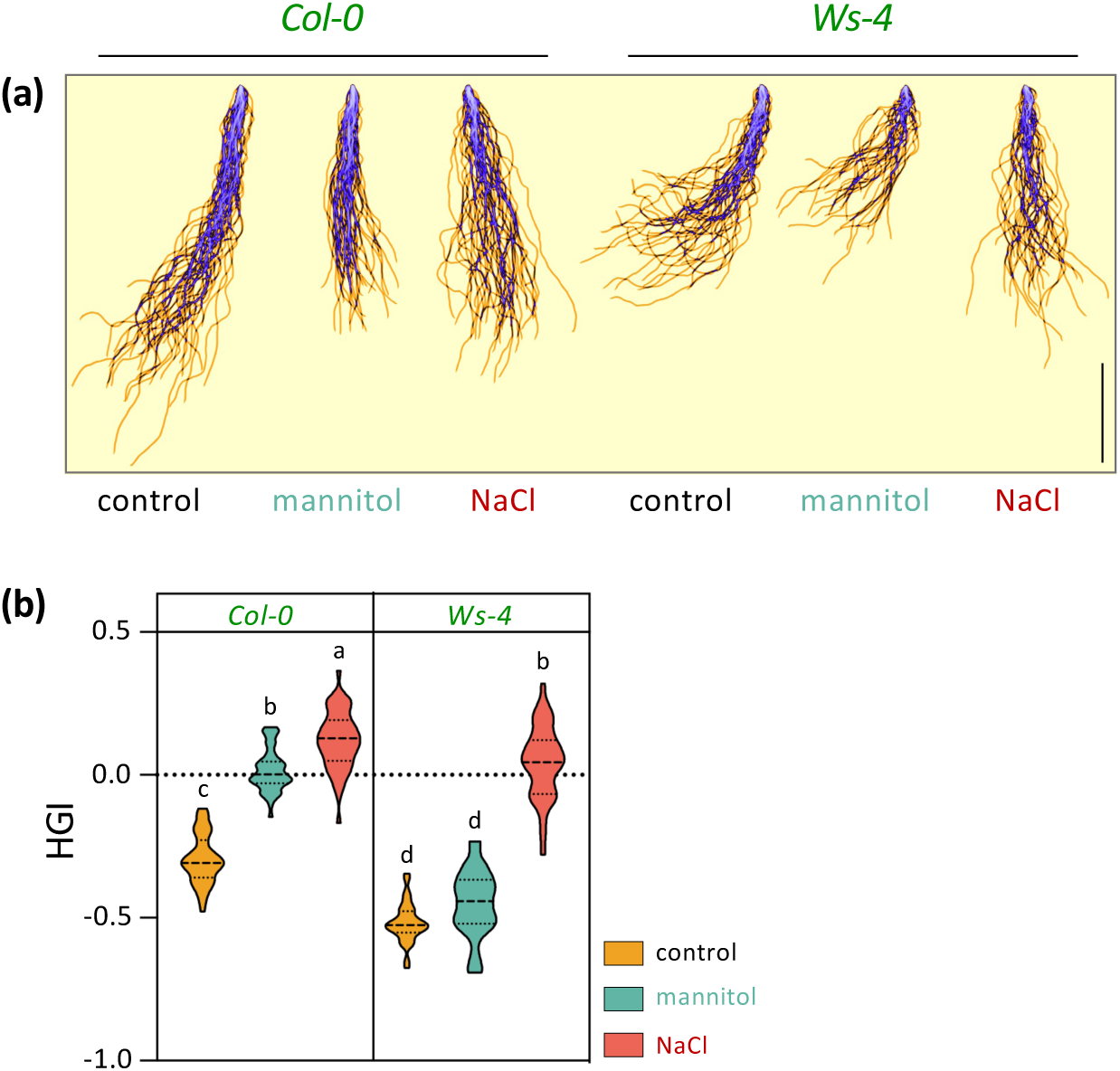
NaCl induced-rightward skewing is mainly ionic but also osmotic. (a) Primary root projections, with overlap in purple, of 9-day-old Col-0 and Ws-4 seedlings, grown on ½MS agar medium containing 800 μM P_i_ with or without 100 mM mannitol or 50 mM NaCl, which have the same osmolality. (b) HGI quantifications. Middle line represents median and dotted line the quartiles. One-way ANOVA was performed to identify significant differences between treatments (*F*_5,236_ = 301.6, *p* < 0.0001). Different letters indicate a statistically significant difference (*p* < 0.05) by Tukey HSD post-hoc analysis (n = 33-45). Scale bar = 1 cm.

To investigate the ionic nature of the reversed skewing response, various Na^+^- and Cl^-^ salts were tested, including KCl, LiCl, NaI and NaBr. For most, a strong toxic effect on root growth was observed (Suppl. Fig. S2), as has been reported by others (Shahzad *et al*., 2016; Shtangeeva *et al*., 2017; Zhang *et al*., 2023). Nonetheless, it was clear that KCl was less potent than NaCl, indicating the cation is most important to induce the rightward root skewing, and that Na^+^ is more effective than K^+^ in both Col-0 and Ws-4 (Suppl. Fig. S2).

### NaCl reverses helical root-growth direction

On vertical agar plates without salt, Arabidopsis roots skew to the left because of the rightward helical movement (circumnutation) of the root tip (Sheng et al., 2024). When agar plates are placed horizontally, roots keep on growing and rotating, which manifests itself as a clockwise (CW) coiling of the root (Fig. 4). To investigate whether NaCl influences helical root growth, seedlings were first grown vertically (70°) for 4 days on ½MS-agar plates containing relatively low (300 µM) or high (2500 µM) P_i_ concentrations, with and without 50 mM NaCl, after which plates were positioned horizontally to follow the helical root growth direction. NaCl reversed the standard CW helical root rotation to counterclockwise (CCW) (Fig. 4a-c), not only in the primary root but also in lateral roots (Fig. 4d, 4e). These results demonstrate that NaCl reverses the helical root-growth direction from CW to CCW in both primary- and lateral roots.

**Fig. 4.**
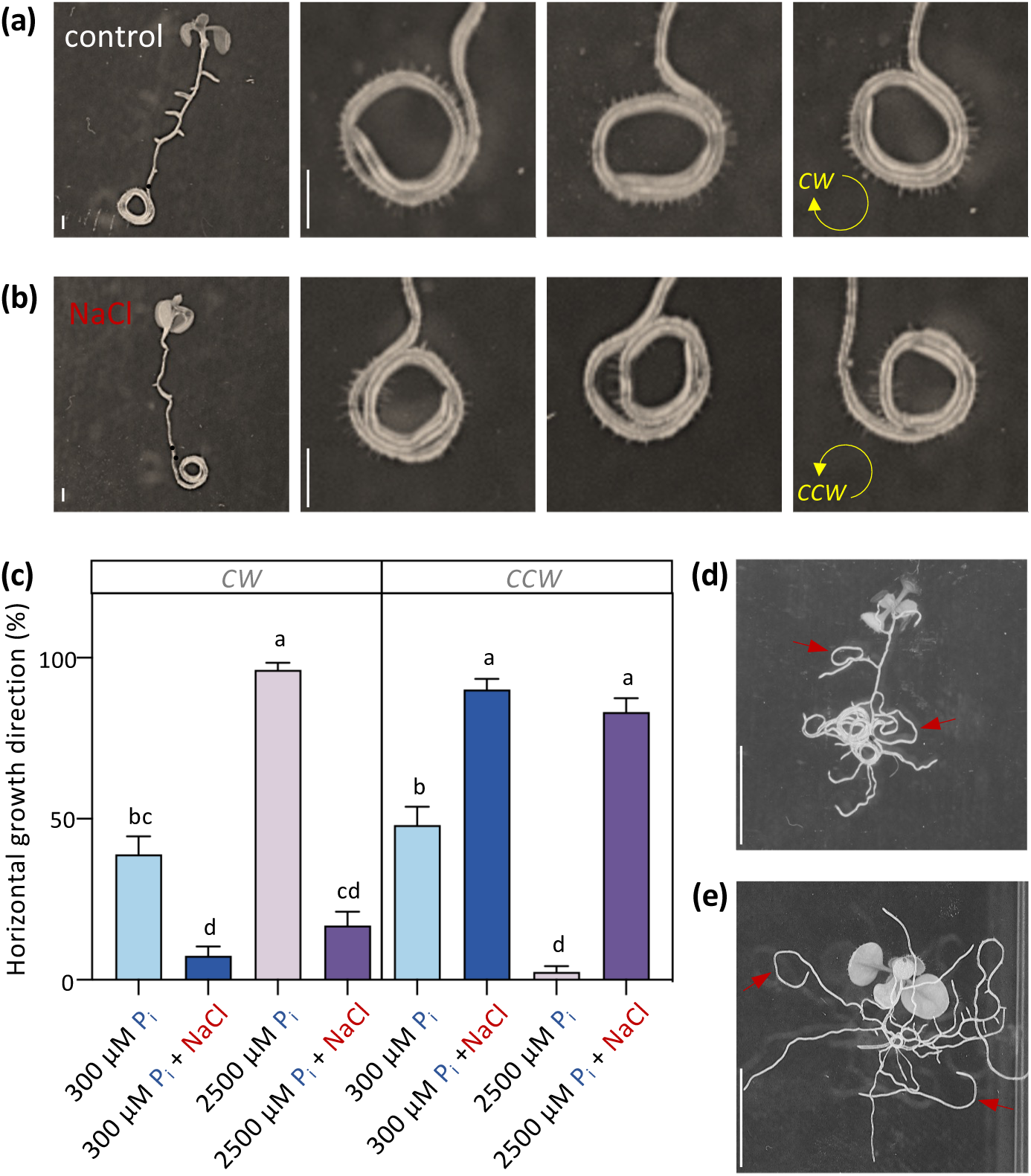
Salinity reverses helical growth direction of Arabidopsis roots. (a, b) Root growth directions in 6-day-old Col-0 seedlings (4 days vertical, 2 days horizontal growth) at 2500 µM P_i_ without (a) or with 50 mM NaCl (b). (c) Quantification of number of roots (%) growing CW or CCW at 300 or 2500 µM P_i_ with and without 50 mM NaCl. (d, e) Helical growth direction in lateral roots growing at 2500 μM P_i_ without-(d) or with 50 mM NaCl (e), after 10 days (4 days vertical, 6 days horizontal) and 14 days (4 days vertical, 10 days horizontal growth), respectively. Red arrows point to CW or CCW curling of lateral roots. Data shown are means ± SEM. Kruskal-Wallis test was performed to identify significant differences. Different letters indicate significant differences (*p*<0.05) among treatments (n = 77-81). Scale bars: (a, b) 1 mm; (d, e) 1 cm.

### NaCl reverses epidermal Cell File Rotation from CCW to CW

Root skewing is closely linked to epidermal CFR in the root elongation zone (Roy & Bassham, 2014; Villaécija-Aguilar *et al*., 2019; Porat *et al*., 2024; Sheng *et al*., 2024). To evaluate the effect of NaCl on CFR, roots of 7-day-old seedlings, grown at 300 or 2500 μM P_i_, and with and without 50 mM NaCl, were examined using stereomicroscopy. At high P_i_, epidermal cell files revealed a pronounced left-handed/CCW rotation (Fig. 5a, b), as we demonstrated before (Sheng *et al*., 2024). In the presence of NaCl, however, an opposite right-handed/CW CFR was observed (Fig. 5a, c). Measuring the angle between root growth and epidermal cell files (Suppl. Fig. S1a) revealed significantly higher CCW angles at high P_i_, and higher CW angles in the presence of NaCl (Fig. 5d). To emphasize the relationship between epidermal CFR and root-growth direction, both phenomena were analysed in roots of seedlings grown on horizontal plates. Without salt, seedlings displayed a right-handed, CW-root growth and left-handed, CCW-CFR (Fig. 5e); *vice versa*, in the presence of NaCl, seedlings exhibited a clear CCW-root growth and CW-CFR (Fig. 5f).

**Fig. 5.**
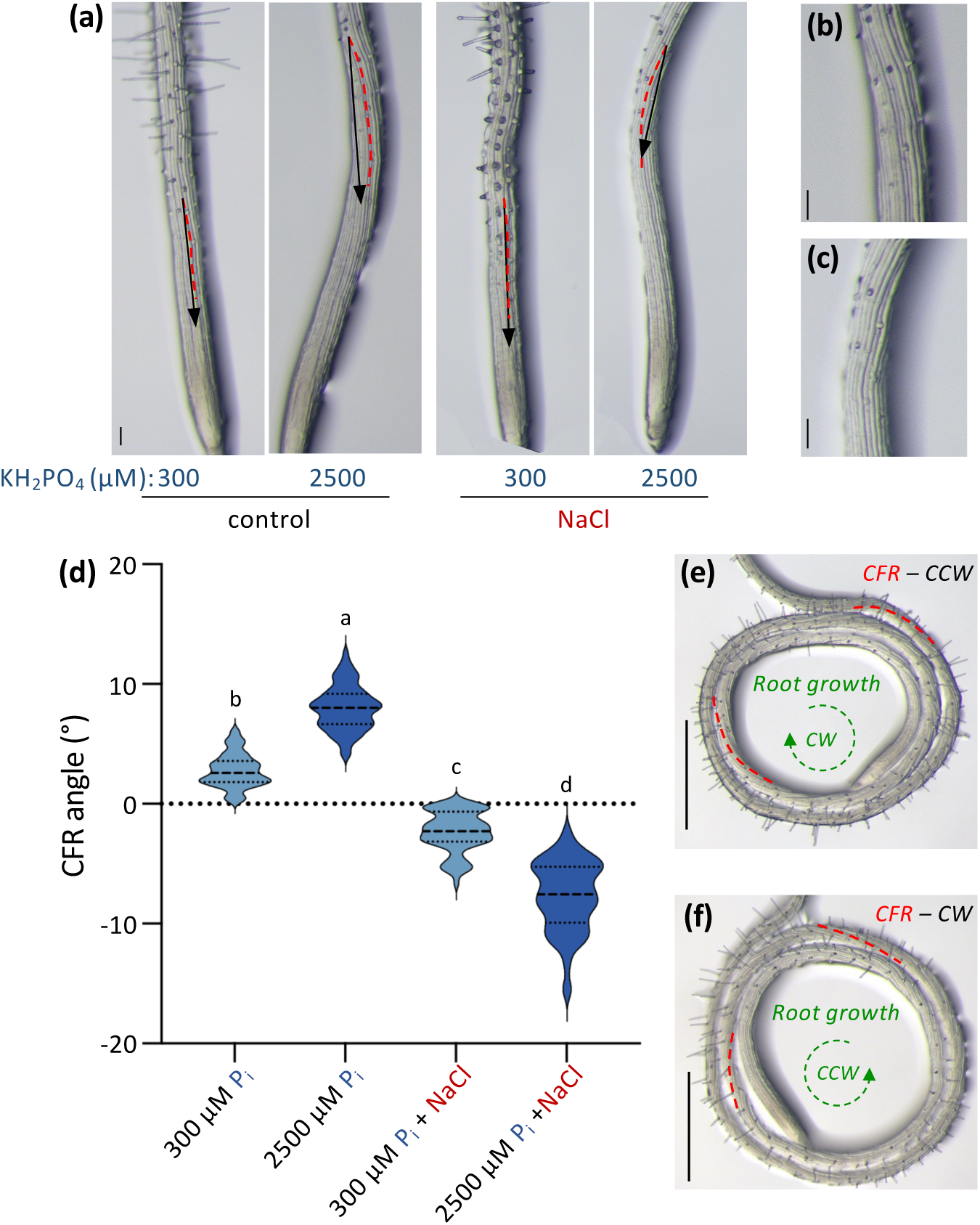
Salinity reverses P_i_-induced epidermal cell-file rotation, from CCW to CW. (a) Stereomicroscopic images of root tips of Col-0 at 300 or 2500 μM P_i_ ± 50 mM NaCl. Black arrows indicate root growth direction; red dotted lines indicate CFR direction. (b, c) Zoom-in of elongation zone without-(b) or with (c) 50 mM NaCl. (d) Quantification of epidermal CFR angles. Significant differences were identified with Kruskal-Wallis test and indicated with different letters (*p* < 0.05) between treatments. Values represent three biological replicates with 13-38 seedlings analysed for each replicate. (e, f) Stereomicroscopic image of horizontal growing Col-0 roots (4 days vertical, 2 days horizontal), showing right-handed (CW) root growth and left-handed (CCW) epidermal CFR on 2500 μM P_i_ medium without NaCl (e), and left-handed (CCW) root growth and right-handed (CW) epidermal CFR with 50 mM NaCl (f). Red dotted line indicates epidermal cell file direction. Scale bars = 1 mm.

Deviations in root skewing and CFR have been related to the organisation of the microtubule cytoskeleton (Ishida *et al*., 2007b; Sedbrook & Kaloriti, 2008; Wang *et al*., 2011b; Buschmann & Borchers, 2020; Sheng *et al*., 2024). Interestingly, salt stress has been shown to affect the organization of the microtubule cytoskeleton (Shoji *et al*., 2006; Wang *et al*., 2007; Zhou *et al*., 2023). To investigate the involvement of microtubules in the salt induced-skewing reversal, we investigated the effects of taxol and propyzamide, which stabilize and destabilize microtubules, respectively (Nakamura *et al*., 2004; Hodge *et al*., 2009). In the presence of taxol, roots showed an enhanced leftward skewing response at both 300 and 2500 µM P_i_, with a stronger effect at high P_i_ (Suppl. Fig. S3; Sheng *et al*., 2024). Similar results were obtained with propyzamide (Suppl. Fig. S4; Sheng *et al*., 2024). In contrast, in the presence of NaCl, taxol reversed the SIRS to the left again in a P_i_-dependent manner, while propyzamide did not (Suppl. Fig. S3 & S4). These results indicate that under salt conditions, P_i_ can only reverse SIRS when microtubules are stabilised. These results are also in agreement with earlier studies, showing that salt stress induces a depolymerisation of cortical microtubules (Wang *et al*., 2007, 2011a; Zhang *et al*., 2012).

### Mutants with altered root skewing

In a GWAS involving early salt-stress responses in root architecture of *Arabidopsis thaliana* seedlings, Deolu-Ajayi *et al*. (2019) identified a number of potential loci, representing 11 candidate genes. To investigate whether these candidate genes may be involved in root skewing, independent homozygous T-DNA insertion lines were obtained and their PDS and SIRS tested (Suppl. Table S3). The majority of the mutants skewed like WT (Col-0) (Suppl. Fig. S5), except independent mutant lines of *GLT1* and *DOB1*. For *dob1-1* and *dob1-2* mutants, increases in both PDS (leftward skewing) and SIRS (rightward skewing) were found (Fig. 6a), with root length being slightly decreased (Fig. 6b). For *GLT1*, all four homozygous T-DNA insertion lines showed a clear decrease in PDS, with *glt1-1* almost completely lacking a PDS response (Fig. 7a, b). In the presence of NaCl, *glt1-1* revealed a hypersensitive SIRS response but this was not observed for any of the other *glt1* alleles, which basically responded like WT (Figs. 7b-e), also when the NaCl concentration was increased further (Suppl. Fig. S6). While findings confirm a role for GLT1 in PDS, its involvement in SIRS requires further investigation. As the T-DNA insertion of *glt1-1* is located at the far 3’-end of the gene (Fig. 7a), and Q-PCR with intermediate primers did produce transcripts (Suppl. Fig. S7), it is possible that *glt1-1* produces a truncated protein, and therefore differs in phenotype compared to the other alleles.

**Fig. 6.**
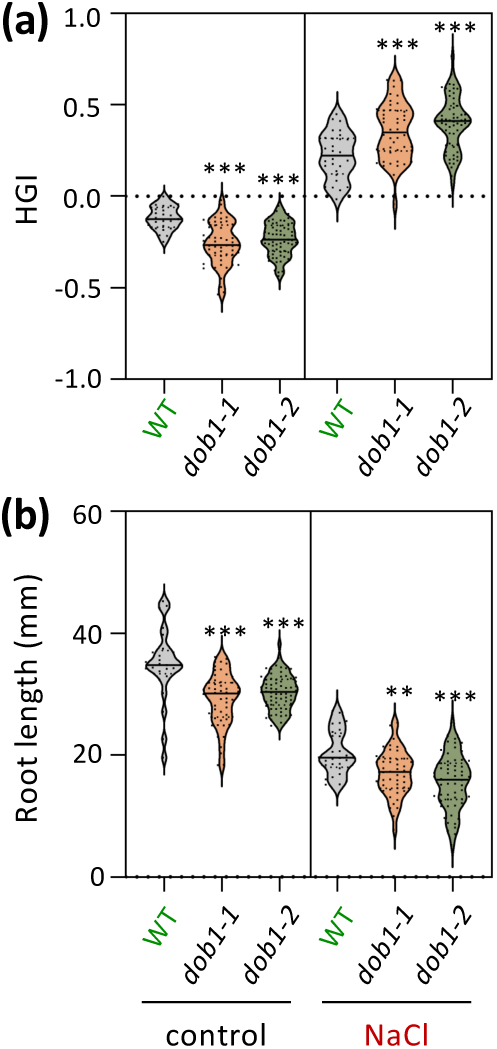
Comparison of root skewing and root length in WT and *dob1* mutants on ½MS media ± NaCl. (a) Quantification of HGI value for WT and mutants grown on ½MS media ± 75 mM NaCl for 9 days. (b) Quantification of root length for WT and mutants. Middle line represents median and dotted line the quartiles. One-way ANOVA was performed to identify significant differences between WT and mutants. Asterisks indicate a statistically significant difference (*P < 0.05, **P < 0.01, ***P < 0.001).

**Fig. 7.**
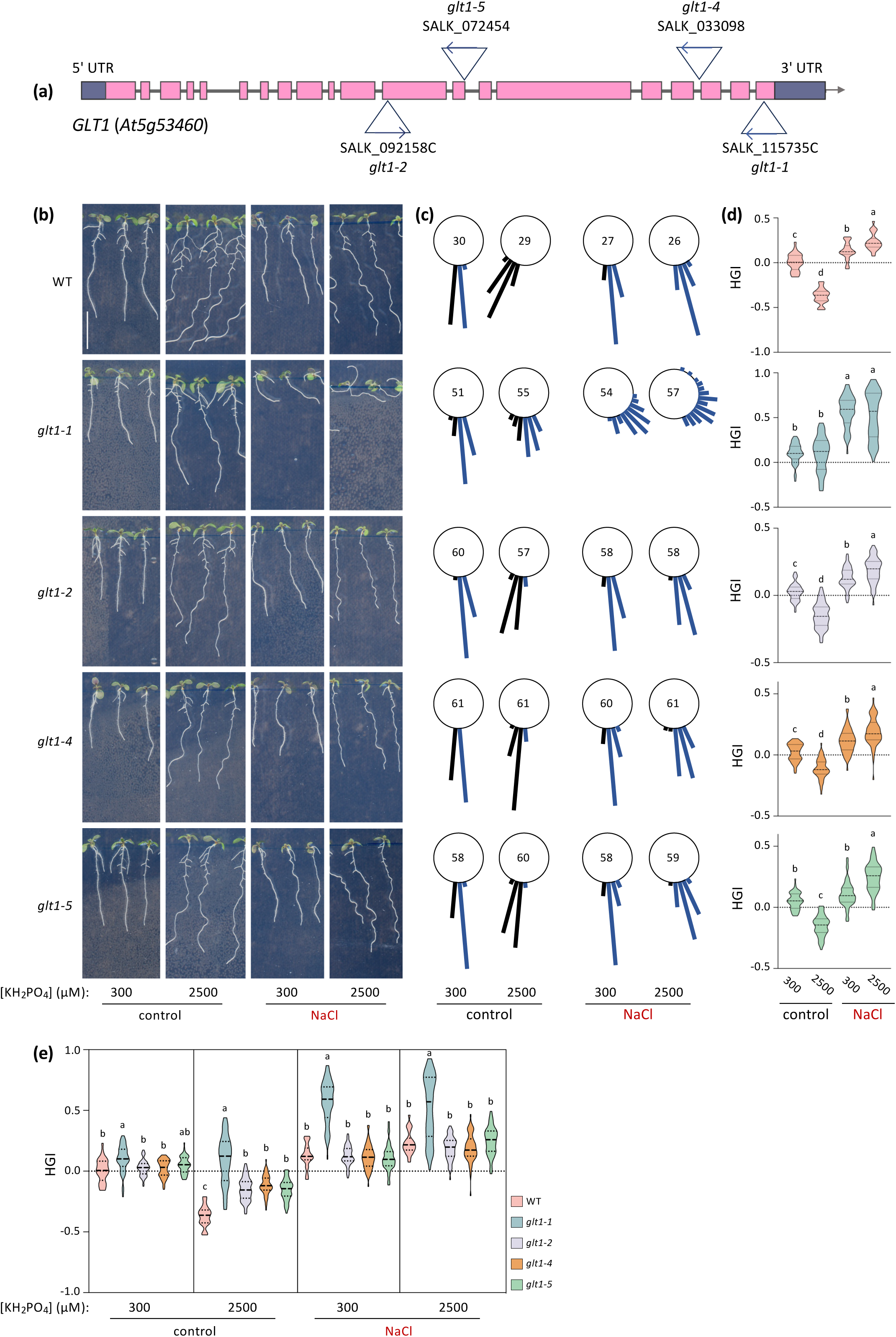
Comparison of root skewing between WT and *glt1* mutants at different P_i_ concentrations ± NaCl. (a) *GLT1* gene indicating positions of T-DNA insertions (triangle). Boxes (pink) and lines indicate exons and introns, respectively. Arrows within triangles indicate the orientation of the left border of the T-DNA insertion. (b) Phenotype of WT and *glt1* mutant seedlings that grown on ½MS agar plates containing either 300- or 2500 µM P_i_ with or without 50 mM NaCl for 9 days. (c) The percentage of roots in angle with the number of roots measured for WT and *glt1* mutant seedlings. (d, e) Quantification of HGI value for WT and *glt1* mutants. Two-way ANOVA was performed to identify significant differences of WT and *glt1* mutants between treatments. Different letters indicate significant differences (*p*<0.05) between treatments. Scale bar = 1cm.

## Discussion

### Salt induced-rightward skewing (SIRS)

Earlier, we showed that Arabidopsis roots skew to the left when grown on vertical ½MS agar plates (seen from the front), and discovered that this is caused by P_i_ in the medium (Sheng *et al*., 2024). This leftward skewing is caused by a rightward/CW rotation of the root tip and a leftward/CCW rotation of epidermal cell files in the root elongation zone. The opposing directional movements are emphasised when seedlings are grown horizontally, showing a clear CW rotational root growth (coiling), coinciding with a CCW (lefthanded) epidermal CFR in the same image (Fig. 8). Using microtubule drugs and FP-based reporters, evidence was provided that the left-handed epidermal CFR, the helical movement of the root tip (circumnutation), and hence the PDS of the root, involves the microtubule cytoskeleton (Sheng *et al*., 2024). Monitoring PDS in various Arabidopsis KO mutants revealed that P_i_-uptake and -signalling are involved, and that PDS is independent of auxin- and strigolactone signalling (Sheng *et al*., 2024).

**Fig. 8.**
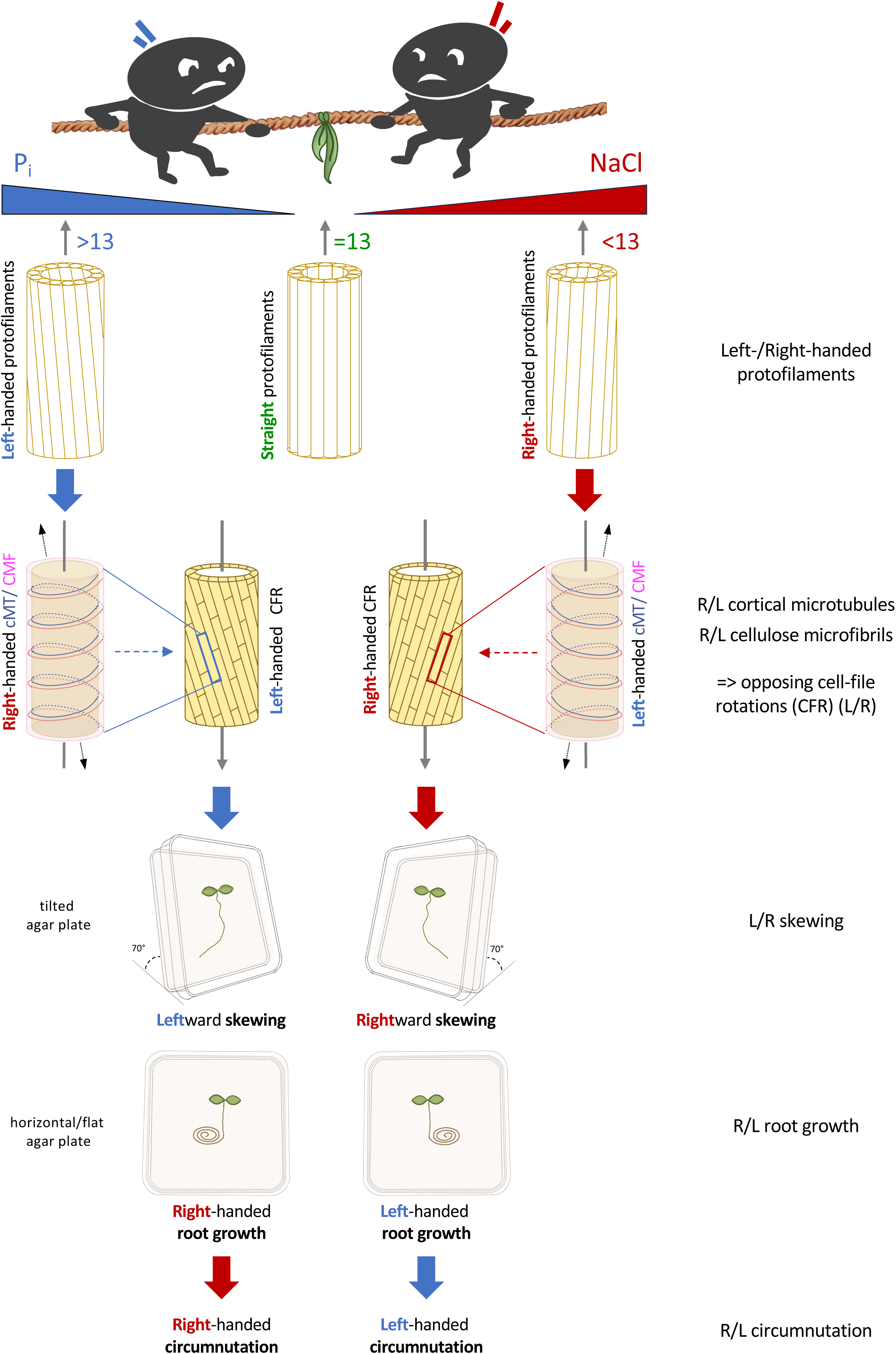
’Tug-of-war’ model, illustrating the opposing effects of P_i_ and NaCl on root skewing based on antagonistic helical movements of various components. The model is based on the helical orientation of cortical microtubules (cMT) due to a change in the number of protofilaments in the microtubule structure and its chirality. Changes impact the direction of the cellulose microfibrils (CMF), the epidermal CFR in the elongation zone and hence, helical growth direction (circumnutation) of the root tip. Increased P_i_ leads to a right-handed shift in cortical microtubules, which causes right-handed twisting of CMFs. The biophysical torsion during cell elongation enhances a left-handed CFR of epidermal cells in the root elongation zone, resulting in clockwise root growth on horizontal agar plates and leftward skewing on vertical plates. NaCl triggers the opposite, a left-handed shift in cMT, causing left-handed twisting of CMFs, which triggers a right-handed CFR that results in a CCW helical root growth and rightward skewing.

In this study, we show that NaCl reverses PDS, by triggering a complete, opposite helical cascade of each of the rotations, ending up with a rightward-skewing response (Figs. 1, 2), like a tug of war, pulling the root in either direction (Fig. 8; Suppl. Table S1). This was again emphasised on horizontal agar plates, where NaCl reversed helical root-growth direction from CW to CCW (Fig. 4) and epidermal CFR from CCW to CW (Fig. 5). While for PDS, relatively big differences between accessions Col-0 and Ws-4 were observed (Sheng *et al*., 2024), for NaCl this is much smaller (Fig. 1-3), indicating that different genetic factors are engaged in driving the reorganisation. The effect of NaCl on root skewing is stronger than of KCl, and much stronger than of mannitol, suggesting the effect is ionic rather than osmotic (Fig. 3, Suppl. Fig. S2). Irrespective, however, our results clearly illustrate that Arabidopsis roots can change their circumnutation, and hence their growth direction in response to ionic- and/or osmotic changes.

### Halotropism, hydrotropism or salt-reversed skewing?

Halotropism represents a relatively new described mechanism for roots to avoid salinity (Galvan-Ampudia *et al*., 2013; Deolu-Ajayi *et al*., 2019; Korver *et al*., 2020; Yu *et al*., 2022; Zheng *et al*., 2024). It was considered a ’tropism’ because of its presumed involvement of auxin (Galvan-Ampudia *et al*., 2013; Korver *et al*., 2020; Zheng *et al*., 2024). The latter was recently disputed, showing halotropism was independent of auxin but was caused by a reorientation of the microtubule cytoskeleton (Yu *et al*., 2022). To measure halotropism, a typical split-diagonal agar system is used, where the left-lower part of the agar medium is replaced by agar medium containing NaCl (*e.g*. 200 mM), which then generates a salt gradient (*e.g*. Deolu-Ajayi *et al*., 2019; Yu *et al*., 2022). As such, roots have always been shown to ’grow away’ from the NaCl gradient. It is, however, troubling that the NaCl source is always placed in the lower left side of the plate, with the consequence that the halotropic root response is always measured as a directional growth response to the right. However, here we show that this is SIRS: independent of any gradient - if there is NaCl in the medium - roots will skew to the right! This means that most, if not all, halotropic responses reported, may actually be SIRS responses. In order to discriminate between halotropism and SIRS, the NaCl source should be offered from the opposite direction, i.e. from the right side of the agar plate. Some halotropism papers appear to have used right-sided NaCl gradients, but that is because these plates have been photographed from the back (and the resulting photos were not turned over) (e.g. Deolu-Ajayi *et al*., 2019). The recently discovered role for microtubules in ’halotropism’ (Yu *et al*., 2022), supports the assumption that halotropism is in fact a SIRS response.

A similar re-analysis may be required for hydrotropism (Eapen *et al*., 2005; Shkolnik *et al*., 2016; Dietrich, 2018). Similar split agar systems have been used for hydrotropism assays but with an osmoticum (sorbitol, mannitol) to generate a difference in water potential. Also there, the left side of the plate seems to be used to provide the ’low water potential’ (osmoticum source), and roots grow to the right, which is interpreted as directionally growing towards a higher water potential. However, reported responses may reflect SIRS too, as shown here for mannitol (Fig. 3, Suppl. Fig. S2). It is imperative that these studies are repeated with the osmoticum source provided from both left and right.

### Role of microtubules in salt induced-rightward skewing

Microtubules are essential for cell division, expansion and morphogenesis, and play a crucial role in enabling plants to adapt to environmental stresses (Onelli *et al*., 2015; Yan *et al*., 2023). Prolonged exposure to salt stress has been shown to trigger a reorganisation of cortical microtubules, including a rapid depolymerisation followed by the reassembly of new MT networks. This dynamic reorganisation increases the plant’s capacity to tolerate and survive saline conditions (Wang *et al*., 2007, 2011a). Salt induced-microtubule depolymerization is also associated with alterations in cytoplasmic Ca^2+^ levels, which in turn favours microtubule reassembly through unknown mechanisms (Wang & Mao, 2019). In previous work, we demonstrated that microtubule drugs that induce MT stabilisation or destabilisation, both promote PDS (Sheng *et al*., 2024). In the present study, the microtubule destabilisation drug, propyzamide enhanced SIRS, while the microtubule stabilising drug, taxol antagonised it (Suppl. Figs. S3&S4). Hence, NaCl-induced depolymerisation of microtubules observed by Wang *et al*. (2007, 2011a), may well drive the SIRS discovered here.

Microtubules are hollow cylindrical tubes consisting of α- and β-tubulin heterodimers (Little & Seehaus, 1988; Cooper, 2000; Elliott & Shaw, 2018), which polymerize in a head-to-tail fashion to form linear polymers, known as protofilaments (Mohri, 1968; Hashimoto, 2015). Typically, microtubules consist of 13 protofilaments, which are aligned in parallel to the microtubule axis, leading to a relatively straight array (Fig. 8) (Chrétien & Wade, 1991; Chrétien *et al*., 1996; Nakajima *et al*., 2004; Pampaloni & Florin, 2008; Chaaban & Brouhard, 2017; Chaaban *et al*., 2018; Nakamura & Hashimoto, 2020). When the number of protofilaments deviates from 13, protofilaments tend to skew along the microtubule axis, forming supertwists (Ishida *et al*., 2007a): with more than 13 protofilaments, left-handed helical twisting results, whereas with less than 13 protofilaments, rightward helical twisting in the microtubule is induced. Since salt is known to reduce the number of protofilaments in microtubules *in vitro* (Böhm *et al*., 1990; Dias & Milligan, 1999), and because NaCl and P_i_ have opposing effects on CFR and root circumnutation, -skewing and -growth, we postulate that the helicity of all these processes are linked to the deviation from 13 protofilaments: a right or left helical turn has inevitable consequences for all helical fates downstream (Fig. 8; Suppl. Table S1). Future analysis, using techniques like cryo-ET (Foster *et al*., 2021) will be required to analyse changes in both number and helicity of protofilaments as a consequence of different P_i_- and NaCl concentrations.

### *Role for GLT1* and *DOB1* in root skewing

A recent GWAS analysis on early root responses to salt stress identified a number of loci that correlated with root system architecture (Deolu-Ajayi *et al*., 2019). Since both PDS and SIRS affect root system architecture, we decided to analyze the 11 genes linked to these loci for their potential involvement in the respective skewing responses using T-DNA insertion mutants (Suppl. Fig. S5). This analysis identified *GLT1* and *DOB1* to play a role in PDS and the latter gene also in SIRS. *GLT1* encodes a glutamate synthetase, which produces glutamate from glutamine and 2-oxoglutarate using NADH (NADH-GOGAT). Glutamate plays an important role in plant growth and development, and in acclimation responses to environmental stresses, including salinity (Qiu *et al*., 2020). What role glutamate or *GLT1* plays in root skewing remains to be elucidated. Potentially, glutamylation of microtubules could play a role, a process that functionalizes disordered C-terminal tubulin tails with glutamate chains of variable lengths. Glutamate chains can be added and removed by tubulin tyrosine ligases (TTLLs) and carboxypeptidases (CCPs) (Bodakuntla *et al*., 2021). In animal cells, its significance in modulating microtubule stability has been well-established but for plants this is still unknown (Torrino *et al*., 2021; Genova *et al*., 2023). Glutamylation negatively regulates tubulin polymerization by reducing growth rates and increasing catastrophe frequencies (Chen & Roll-Mecak, 2023).

Another possibility is that it is related to touch sensing. In a recent study, touch was suggested to trigger the release of glutamate, which diffuses locally through the apoplast, activating the calcium-permeable channel, GLUTAMATE RECEPTOR-LIKE 3.3 (Bellandi *et al*., 2022). Interestingly, the release of glutamate upon touch was proposed to influence microtubule dynamics and properties via a post-translational modification of tubulin. The reported association between glutamate and microtubules in combination with the *glt1* phenotype may suggest a functional relationship and deserves further study in plants.

*DOB1* is an unknown gene that is upregulated in roots during salt stress, and was recently suggested to be involved in halotropism, since T-DNA insertion mutants exhibited reduced halotropic root responses in Arabidopsis (Deolu-Ajayi *et al*., 2019). In the present study, we demonstrated that *dob1-1* and *dob1-2* exhibit enhanced PDS. With salt, however, the two *dob1* mutants showed opposite results, with *dob1-1* showing reduced and *dob1-2* increased SIRS (Fig. 6). Since both *dob1* mutants were reported KO mutants (Deolu-Ajayi *et al*., 2019), the different phenotypes observed in the presence of NaCl might be due to differences in the effect of the T-DNA insertion site on protein binding or regulatory control and requires further exploration in the future.

### Physiological consequences

The circumnutating movement, as described by Darwin (Darwin & Darwin, 1880), is regarded as a fundamental mechanism from which all other movements may have evolved. Such movement can occur in either left-handed (CCW) or right-handed (CW) direction (Migliaccio *et al*., 2013). Recent research on rice roots showed circumnutation in both directions, and that this was crucial for roots to penetrate soil and to circumvent obstacles (Taylor *et al*., 2021). In Arabidopsis, right-handed (CW) helical root growth (and leftward skewing) appears to be dominant (Simmons *et al*., 1995; Sheng *et al*., 2024). Mutants that skew rightward and exhibit CCW helical growth, like *spiral* mutants, *spr1* and *spr2*/*tor1*, turned out to encode plant-specific microtubule-associated proteins that regulate the orientation of cortical microtubules and the direction of organ growth (Furutani *et al*., 2000; Nakajima *et al*., 2004; Buschmann *et al*., 2004). An increase in root skewing by P_i_ may help to keep the root system in the topsoil where P_i_ tends to accumulate because of its strong negative charge (Sheng *et al*., 2024). Interestingly, this is a ’force’ that overrules the gravitropic root response that normally pulls the root down. Given the importance of P_i_ for plant growth and development throughout their life cycle, mechanistically this could be driven by evolution.

It is tempting to speculate that SIRS evolved similarly, generating a shallower root system that would be beneficial to escape salinity as cations are washed out easier than anions. What should be kept in mind, however, is that glycophytes did not typically evolve in the presence of NaCl, which primarily arose from agriculture and irrigation. It will be crucial to see whether PDS, and its reversal by salt, is manifested in other plant species. Interestingly, however, in rice, a shallow root system has been found beneficial for both P_i_ uptake (Oo *et al*., 2021) and salt tolerance (Kitomi *et al*., 2020) compared to deeper rooting system.

So far, circumnutation has not been associated to P_i_ nor salt. In maize, root circumnutation is more pronounced under light, while in darkness, roots tend to grow towards the gravity vector (Yokawa & Baluška, 2018). In rice, root circumnutation was induced by high ambient temperature and dependent on ethylene (Cai *et al*., 2024). In pea, root circumnutation has been coupled to gravitropic processes via auxin (Kim *et al*., 2016).

Whether a root turns left or right, this may in the end not be that important, as long as it turns, since this will facilitate soil penetration and help to circumvent obstacles. Increased circumnutation may help, however, to grow a more shallow root system architecture and to explore the soil more efficiently. Drought and gravity may overrule such responses again. A better understanding of the root’s potential to twist and rotate to improve root system architecture may present breeders with novel tools to breed for crops that explore soil environments better, thereby contributing to improved agricultural practices and crop productivity.

## Author Contributions

The project was conceived and experiments were designed by HS, HB and TM. Most experimental work and analyses were performed by HS. Gene expression analysis was performed by RvW. HS and TM constructed figures and wrote the manuscript. All authors read and approved the final manuscript.

## Data availability

All data generated and/or analysed during this study are included in this paper and its Supplementary Information files. Materials used in this study are available from the corresponding author upon reasonable request. Source data are provided within the paper.

