## Supplemental files for "Salt stress reverses root circumnutation, -skewing and -growth direction in Arabidopsis"

### Supplemental Figures

Sheng et al., 2026

| | helicial turn/rotational direction | |
|---|---|---|
| **phenotype** | **$P_i$** | **NaCl** |
| horizontal root growth | right/CW | left/CCW |
| circumnutation | right | left |
| skewing | left | right |
| cell file rotation (CFR) | left | right |
| cortical microtubules (cMT) | right | left |
| cellulose microfibrils (CMF) | right | left |
| protofilaments | left | right |
| nr. of protofilaments | >13 | <13 |

Supplemental Table S1

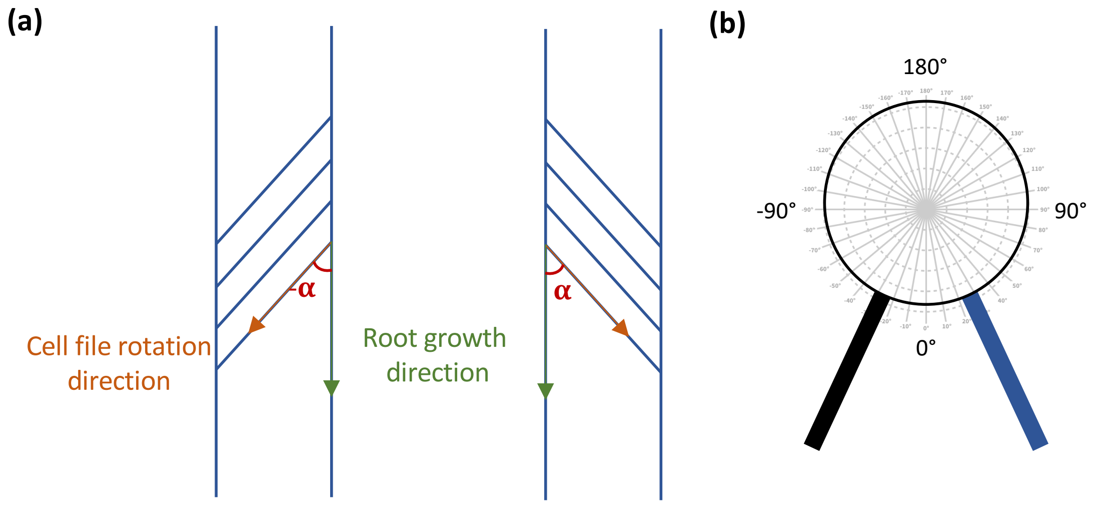


Supplemental Figure S1

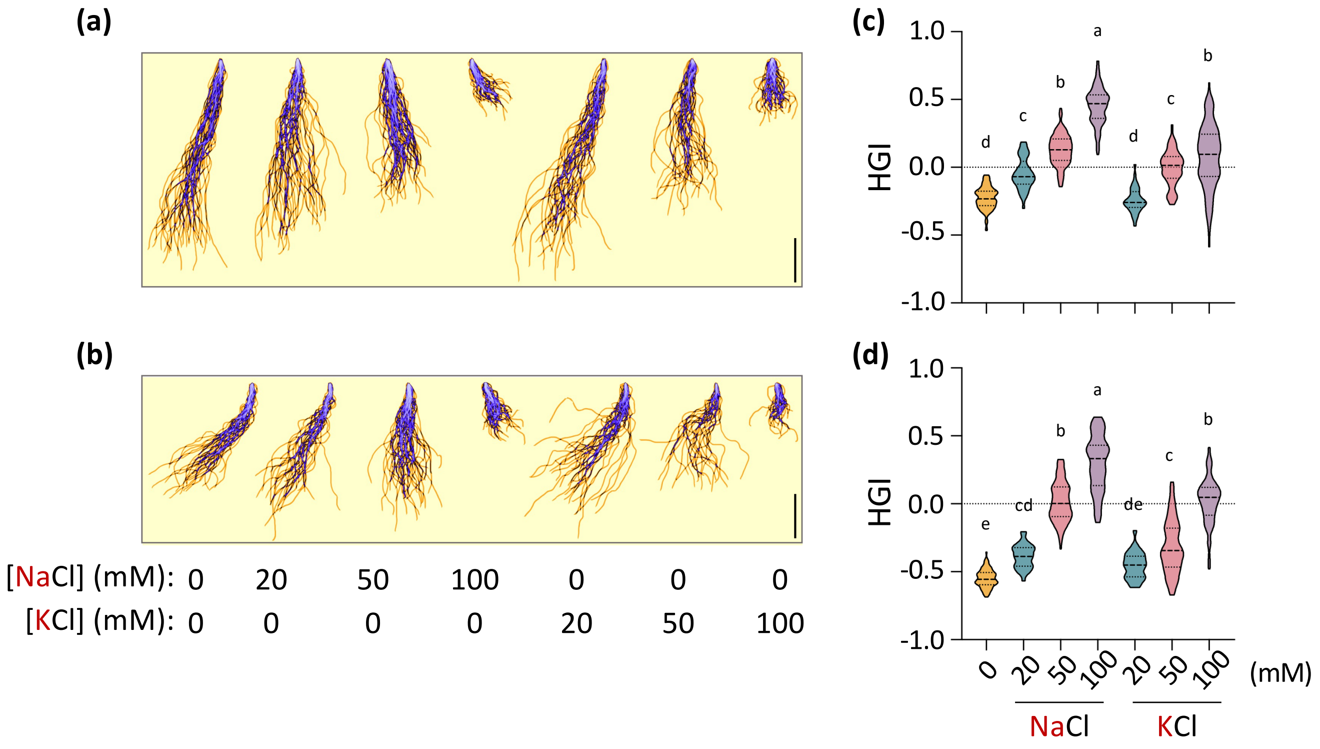


Supplemental Figure S2

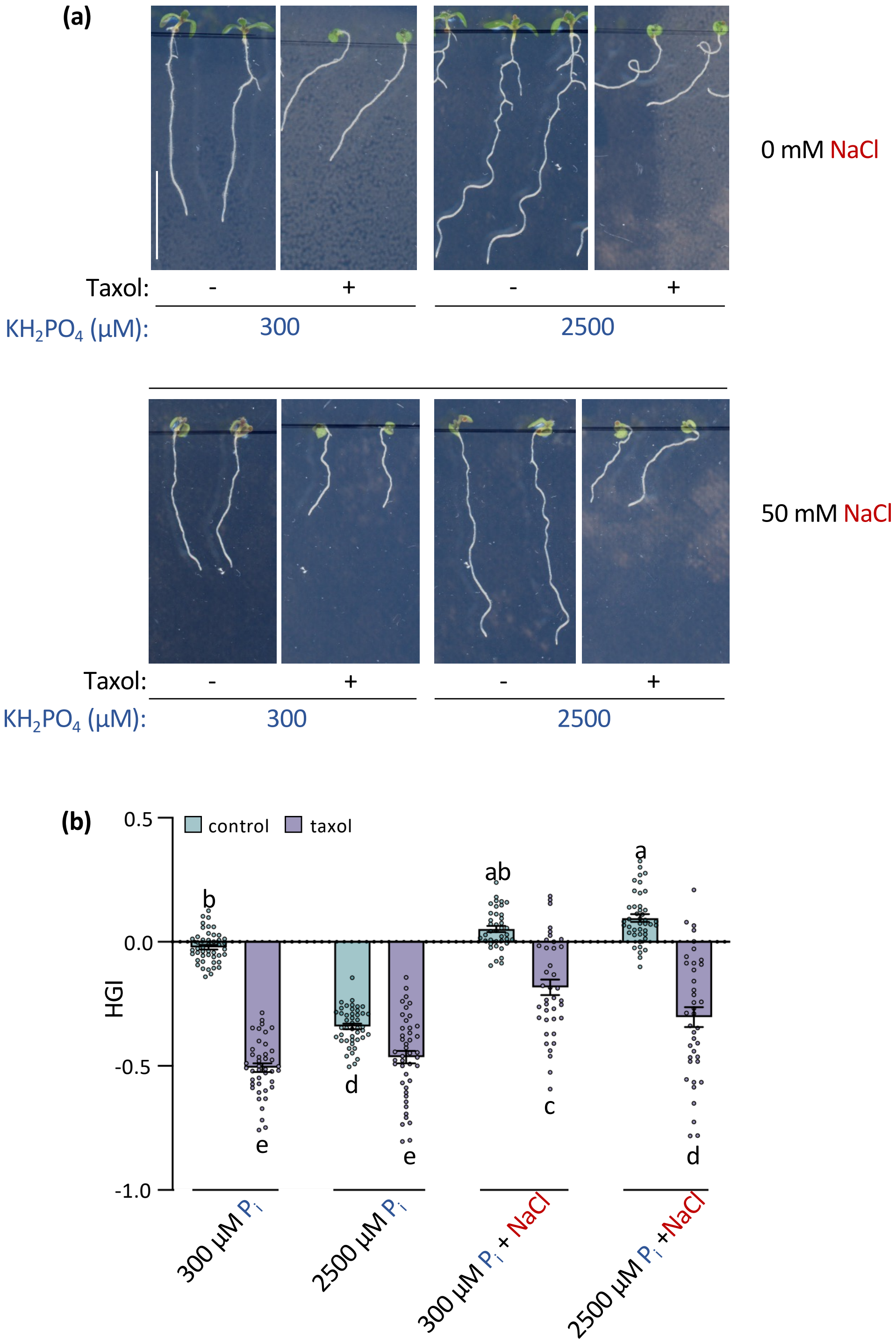


Supplemental Figure S3

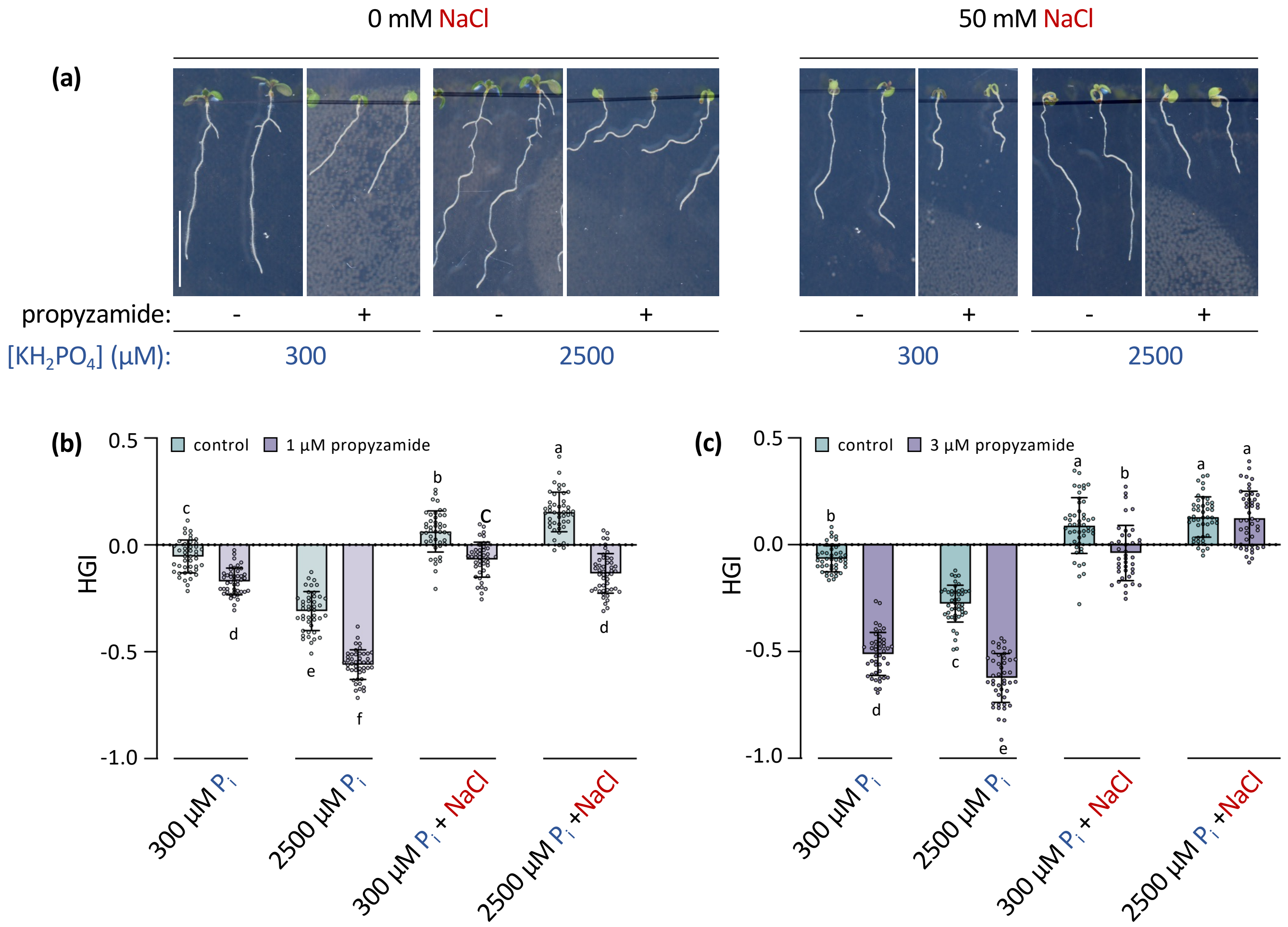


Supplemental Figure S4

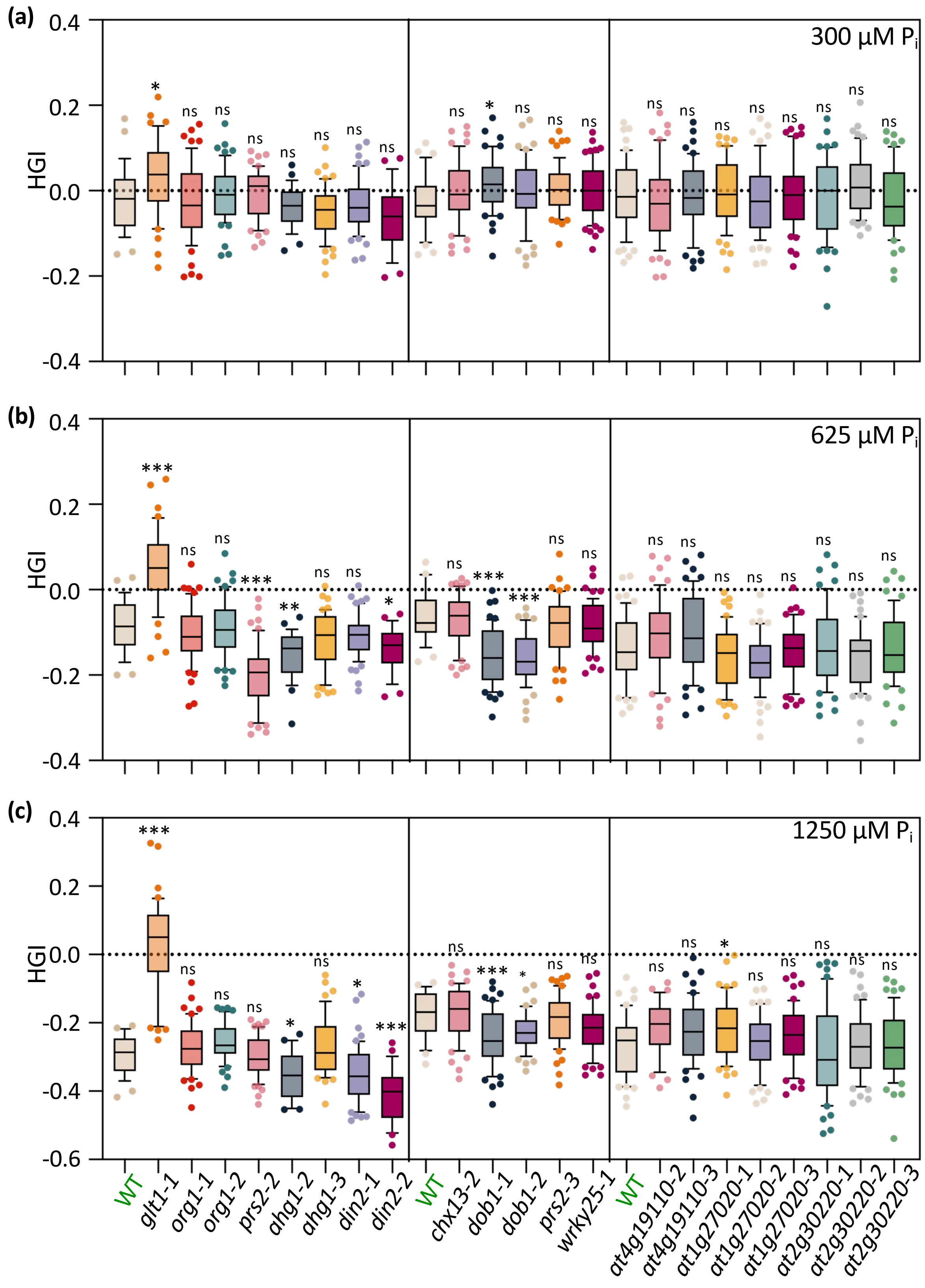


Supplemental Figure S5

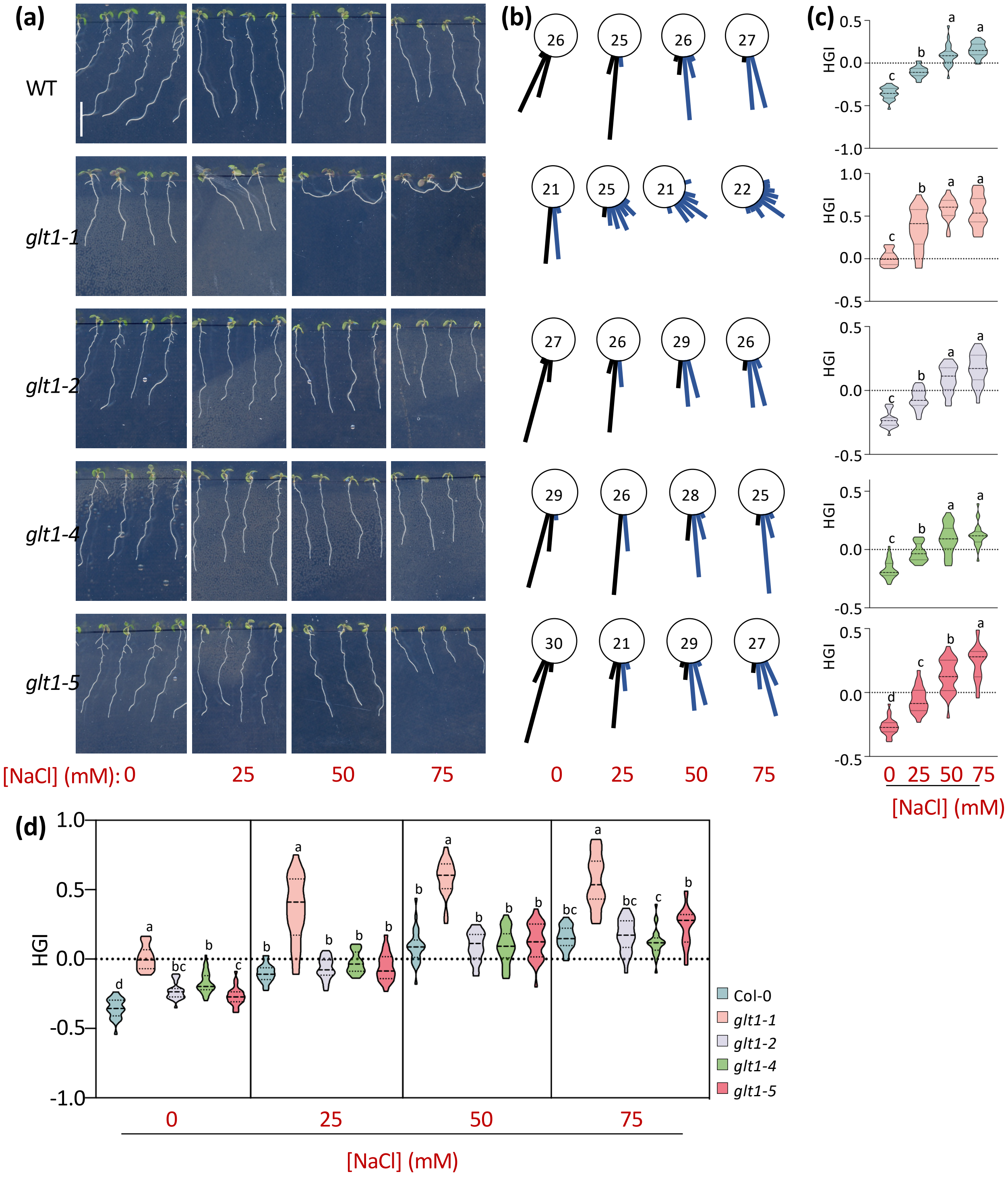


Supplemental Figure S6

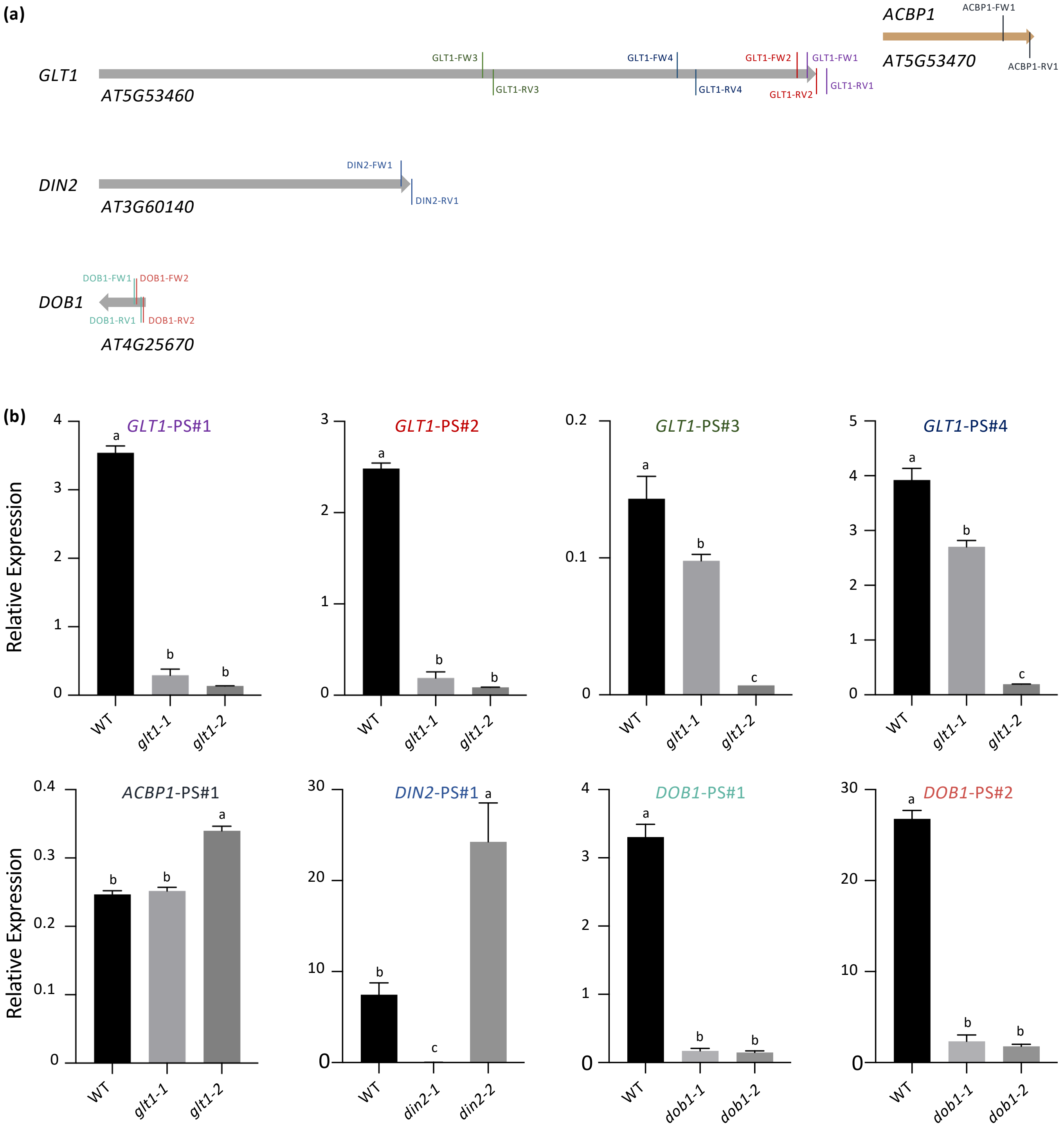


Supplemental Figure S7
